# A Synthetic Dynamic Polyvinyl Alcohol Photoresin for Fast Volumetric Bioprinting of Functional Ultrasoft Hydrogel Constructs

**DOI:** 10.1101/2022.10.20.513079

**Authors:** Wanwan Qiu, Jenny Gehlen, Margherita Bernero, Christian Gehre, Gian Nutal Schädli, Ralph Müller, Xiao-Hua Qin

**Affiliations:** Institute for Biomechanics, ETH Zürich, Leopold-Ruzicka-Weg 4, 8093 Zürich, Switzerland

**Keywords:** volumetric bioprinting, hydrogels, bioresins, polyvinyl alcohol, thiol-ene reaction

## Abstract

Tomographic volumetric bioprinting (VBP) enables fast photofabrication of cell-laden hydrogel constructs in one step, addressing the limitations of conventional layer-by-layer additive manufacturing. However, existing biomaterials that fulfill the physicochemical requirements of VBP are limited to gelatin-based photoresins of high polymer concentrations. The printed microenvironments are predominantly static and stiff, lacking sufficient capacity to support 3D cell growth. We here report a dynamic resin based on thiol-ene photo-clickable polyvinyl alcohol (PVA) and thermo-sensitive sacrificial gelatin for fast VBP of functional ultrasoft cell-laden hydrogel constructs within 7-15 seconds. Using gelatin allows VBP of permissive hydrogels with low PVA contents of 1.5%, providing a stress-relaxing environment for fast cell spreading, 3D osteogenic differentiation of embedded human mesenchymal stem cells and matrix mineralization. Additionally, site-specific immobilization of molecules-of-interest inside a PVA hydrogel is achieved by 4D tomographic thiol-ene photopatterning. This technique may enable spatiotemporal control of cell-material interactions and guided *in vitro* tissue formation using programmed cell-friendly light. Altogether, this study introduces a synthetic dynamic photoresin enabling fast VBP of functional ultrasoft hydrogel constructs with well-defined physicochemical properties and high efficiency.

## 1. Introduction

Recent advances in three-dimensional (3D) bioprinting^[1–4]^ have witnessed the fabrication of various 3D tissue models for both fundamental and translational research in regenerative medicine. Conventional 3D bioprinting techniques rely on the extrusion of a shear-thinning bioink through a nozzle causing high shear stress which can compromise the integrity of embedded stem cells.^[5]^ Layer-by-layer fabrication is a slow process with limited possibilities in structural design. Fabrication of soft hydrogels containing a permissive tubular structure for example remains difficult because the substrates tend to deform when new layers are added.

To address these limitations, volumetric bioprinting (VBP) has recently emerged as an enabling technique that utilizes tomographic back-projections to fabricate centimeter-scale cell-laden 3D hydrogel constructs within tens of seconds.^[6–8]^ In VBP, a 3D object is created in a single step by irradiating a photoresin from multiple angles (**Figure 1a**). The computed light patterns, which are generated by a digital light projector (DLP) modulator, are spatially projected into a rotating glass vial containing the resin. Accumulation of light dose induces localized solidification of the resin through crosslinking reactions. The rotational motion during VBP requires high viscosity resins to stabilize the formed structures.

**Figure 1.**
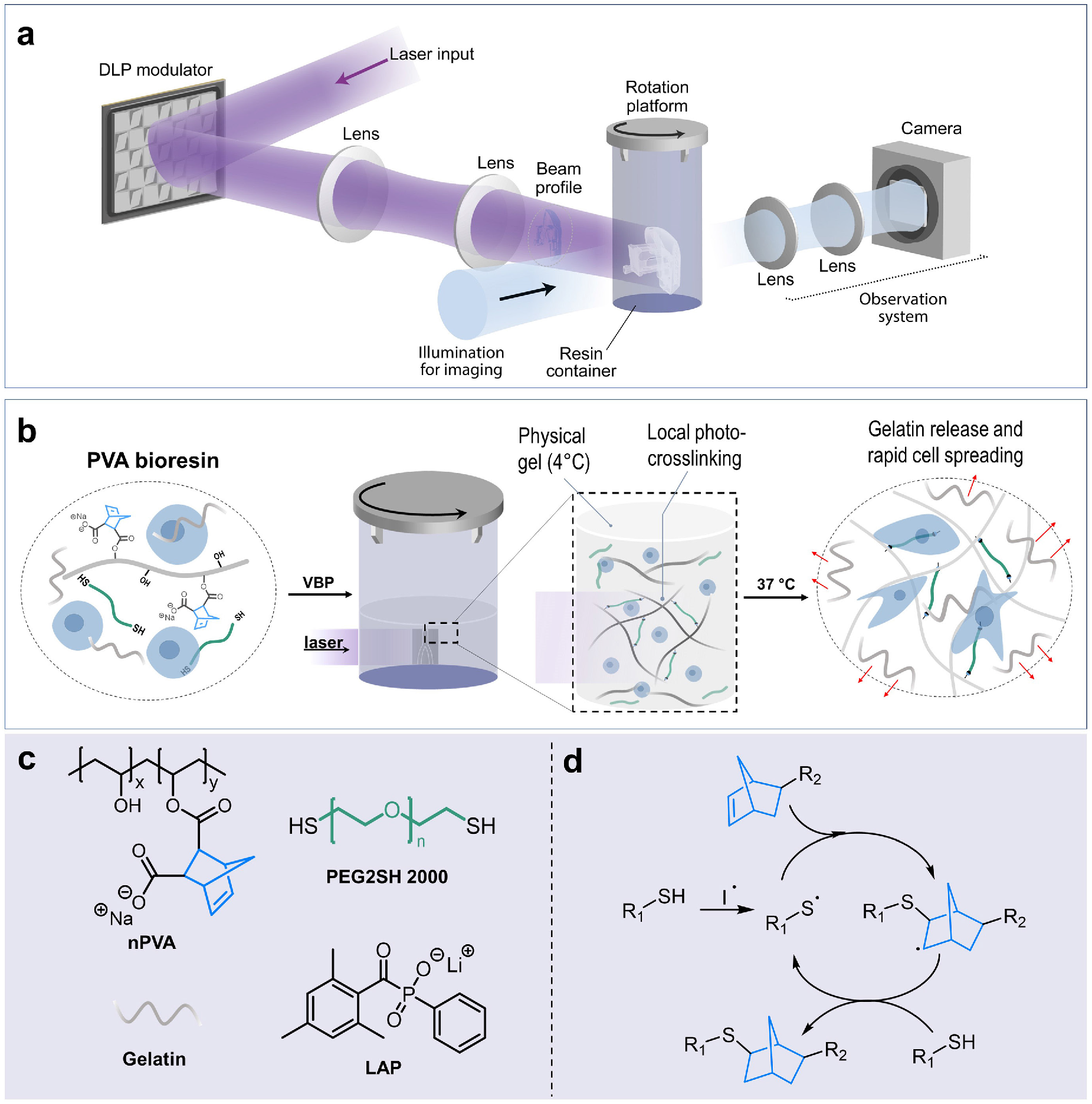
Design of a synthetic dynamic photo-clickable resin for VBP. (**a**) Schematic of the set-up for VBP. Reproduced under the terms of the CC-BY Creative Commons Attribution 4.0 International license. Copyright 2020, Nature Communications.^[7]^ (**b**) Illustration of VBP of a PVA bioresin following three steps: cooling-induced solidification of the resin, tomographic photocrosslinking, and thermal liberation of gelatin at 37°C after printing. (**c**) Chemical structures of nPVA, thiolated crosslinker (PEG2SH) and photoinitiator (LAP). (**d**) Mechanism of radical-mediated thiol-norbornene photo-click reaction: the activated initiator generates free radical that abstracts a hydrogen atom from a thiol to give a thiyl radical which propagates across the norbornene C=C bond. The produced norbornane radical abstracts a hydrogen from the thiol group to regenerate a thiyl radical and form a thioether bond.

Since the first report on VBP in 2019,^[6]^ there has been an increasing interest in developing new bioresins for VBP.^[9–10]^ For instance, Bernal et al.^[6]^ reported a bioresin of 10% gelatin methacryloyl (GelMA) and 0.037% (1.26 mmol m^−3^) Lithium phenyl-2,4,6-trimethylbenzoylphosphinate (LAP) as photoinitiator. By combining VBP and subsequent photochemical post-curing, stable cell-laden hydrogel structures were fabricated. Thereby an increased polymer concentration in the resin improved the printability but the higher stiffness of the printed matrices proved to be less suitable to support 3D cell growth. Recently, Gehlen et al. developed a soft bioresin composed of 5% GelMA and 0.05% LAP (1.7 mmol m^−3^) which showed enhanced *in vitro* osteogenic differentiation of human mesenchymal stem cells (hMSC) when compared to a control experiment using 10% GelMA.^[10]^ Rizzo et al.^[9]^ reported a thiol-ene resin based on norbornene-functionalized gelatin^[11–13]^ and 4-arm poly(ethylene glycol) thiols. Using this resin, free-form hydrogel constructs containing C2C12 myoblasts were printed with good stability. All these reported bioresins are based on photocurable gelatin derivatives and use high polymer concentrations. The printed matrices therefore show high mechanical stiffness which limits fast cell spreading and 3D cell growth. Using these hydrogels makes it difficult to control different properties of the matrix such as stiffness and adhesiveness independently. Furthermore, limitation to proteinaceous hydrogels makes a comparison of cell-matrix interactions on a mechanistic level with those observed in synthetic hydrogels (PEG,^[14–18]^ polyvinyl alcohol,^[19–20]^ and polyacrylamide^[21–22]^) difficult. These limitations motivated us to develop a general strategy to expand the choice of bioresins applied in VBP and to develop a low polymer concentration resin by combining norbornene functionalized polyvinyl alcohol (nPVA) and thiolated PEG with gelatin as sacrificial component. We show that combining nPVA with gelatin ensures good printability during the VBP process, while thermal removal of gelatin after the printing process enables PVA-based matrices with low stiffness for efficient 3D cell growth.

We prepared a series of bioresins with varying nPVA content and demonstrate that physically crosslinked gels are formed upon cooling to 4°C (Figure 1b). These bioresins rapidly photocure via highly efficient radical-mediated thiol-ene click chemistry (Figure 1c). Impressively VBP of complex permissive hydrogel constructs within 7-15 s at extremely low nPVA content of only to 1.5% can be realized. Additionally, the printed matrices exhibit a stress-relaxing behavior, which supports fast cell spreading already 2 h after VBP and the formation of 3D bone cellular networks after 14 days of osteogenic differentiation in a soft matrix. Finally, we demonstrate fast site-specific immobilization of molecule-of-interest within a hydrogel by 4D tomographic thiol-ene photopatterning, which is currently unachieved in conventional photoresins.^[6, 23–25]^

## 2. Results and Discussion

### 2.1. Design of a Synthetic Dynamic Photoresin for VBP

Resins applied in VBP need to have a high viscosity, good cell-compatibility and high photocrosslinking efficiency. Recently, Ouyang et al.^[26]^ reported a universal strategy to mix gelatin with a variety of inks (alginate, PEG acrylates, hyaluronan, GelMA) for extrusion bioprinting. Based on these results we therefore decided in favor of a formulation containing a photocrosslinkable synthetic network which provides a modular 3D environment to embedded cells and a sacrificial thermo-reversible network which enables printability. The photocrosslinkable synthetic network was realized by an off-stoichiometric mixture of nPVA and PEG di-thiol (thiol:ene = 4:5) to ensure a facile synthesis of a low cost covalent network by highly reactive thiol-ene photocrosslinking.^[20, 27–28]^ We reason that the incorporation of sacrificial gelatin in this synthetic hydrogel and its subsequent thermal removal at 37°C will greatly expand the scope of resins suitable for VBP and render the printed environments more permissive for 3D cell growth.

Our printing method relies on three key steps: cooling-induced gelation of the resin, tomographic photocrosslinking using computer-aided design (CAD) models, and eventually thermal removal of gelatin at 37°C (Figure 1b). Additionally, non-reacted norbornene moieties are present after photocuring, which can be used for biofunctionalization using a cysteine-containing fibronectin-derived RGD peptide (CGRGDS) to promote cell attachment. LAP^[29]^ was selected as the photoinitiator because of its cytocompatibility and high initiation efficiency. Using this design principle we can independently control the printability and physicochemical properties of the resin (**Table S1**), which allows us to create dynamic hydrogel environments.

### 2.2. Rheological and Physical Properties of PVA Hydrogels

We first investigated the effect of gelatin addition on the rheological properties by a temperature sweep experiment between 37°C and 4°C using a resin composed of 2% nPVA (Mix 3, Table 1) and 5% gelatin. The resin showed a complex modulus of ~3 Pa at 37°C and sol-gel transition at ~18°C. When cooling to 4°C the temperature-dependent complex modulus reached 1000 Pa where the formation of a physical gel was observed (storage modulus G’ >> loss modulus G’’) (**Figure 2a**). This is a first sign that the resin can be applied in VBP. Looking at the rheological properties of a resin containing 2% nPVA without gelatin we observed a complex modulus of only ~1 Pa during the temperature sweep experiment. From these results, we infer that the inclusion of gelatin indeed is important to ensure the printability of low polymer concentration resins.

**Table 1.** Physicochemical properties of PVA hydrogels with varying polymer concentration.

| Group <sup>a)</sup> | Resin <sup>c)</sup> | Storage modulus<br>G' (kPa) <sup>d)</sup> | Loss modulus G''<br>(Pa) <sup>d)</sup> | Loss factor<br>($\times 10^{-3}$) <sup>d)</sup> | Compressive modulus E<br>(kPa) <sup>e)</sup> |
| --- | --- | --- | --- | --- | --- |
| Mix 1 <sup>b)</sup> | 1% nPVA | 3.4 ± 0.4 | 32.0 ± 3.9 | 9.5 ± 0.3 | - |
| Mix 2 | 1.5% nPVA | 6.6 ± 0.2 | 56.9 ± 1.4 | 8.6 ± 0.3 | 2.7 ± 0.1 |
| Mix 3 | 2% nPVA | 7.5 ± 1.3 | 56.6 ± 9.4 | 7.6 ± 0.2 | 6.6 ± 0.4 |
| Mix 4 | 3% nPVA | 24.0 ± 3.5 | 131.4 ± 44.4 | 5.4 ± 1.7 | 25.1 ± 1.7 |
| Mix 5 | 10% GelMA<br>(reference) | 59.7 ± 1.2 | 954.7 ± 22.9 | 16.0 ± 0.2 | 139.0 ± 6.7 |

<sup>a)</sup> All resins were supplemented with 5% gelatin except Mix 5; <sup>b)</sup> Mix 1 cannot form a stable enough gel; <sup>c)</sup> thiol:ene = 4:5; <sup>d)</sup> Plateau values after *in situ* photocrosslinking for 300 s; <sup>e)</sup> Measured after 7 days incubation in PBS at 37°C.

**Figure 2.**
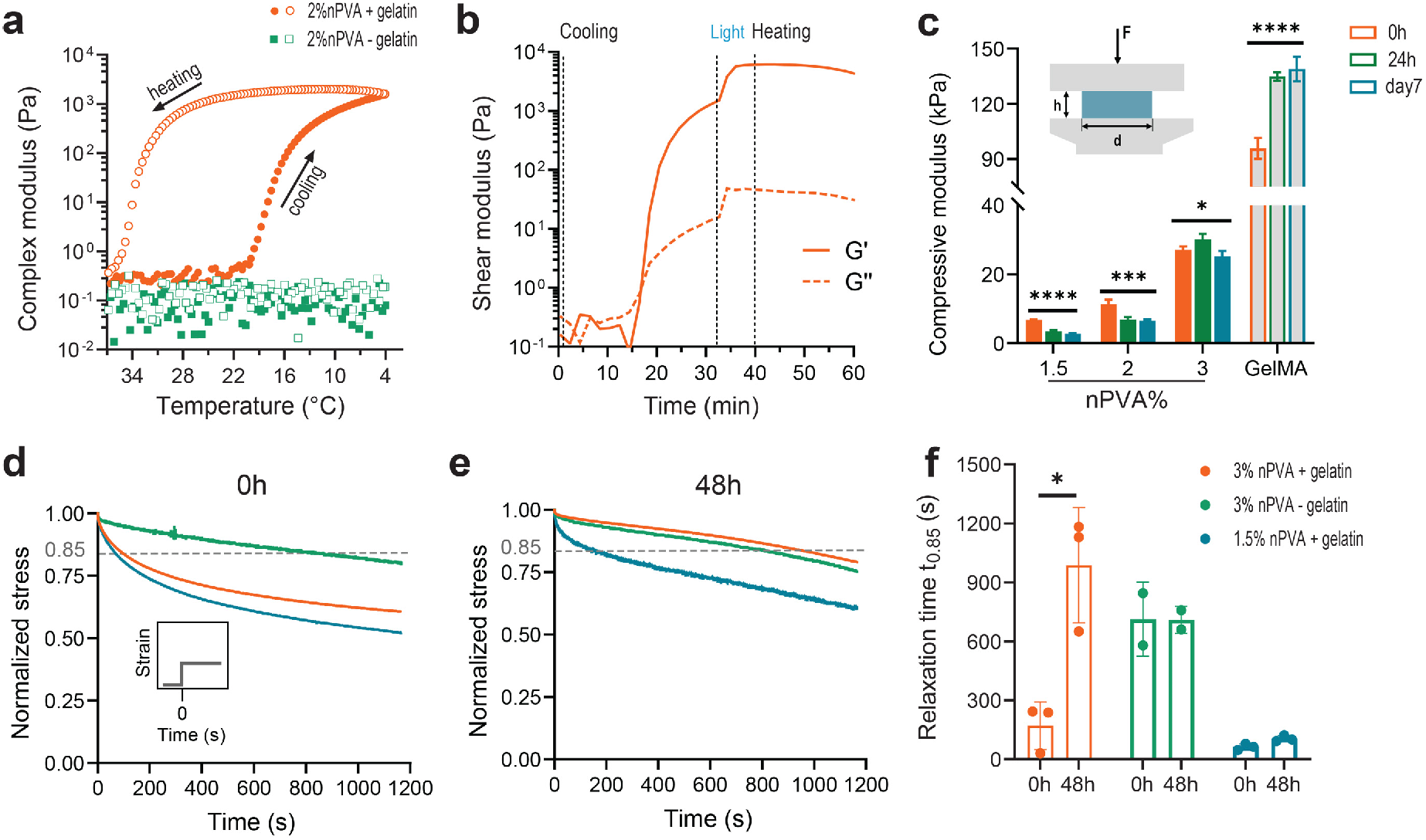
Rheological and mechanical properties of PVA hydrogels. (**a**) Oscillatory temperature sweep (between 37°C and 4 °C at a rate of 1 °C min^−1^; closed symbols denote the cooling whereas open ones denote the heating) showing thermo-reversible gelation of a nPVA/gelatin resin, while an undoped 2% nPVA resin exhibits a low viscosity across the range. (**b**) Storage (G’) and loss (G’’) moduli during a time sweep of a 2% nPVA resin during stages of cooling (from 37°C to 4°C, 1°C min^−1^), UV light exposure (365 nm, 10 mW cm^−2^), and heating back to 37°C. **(c)**The compressive moduli (*E*) of PVA hydrogel disks before (0 day) and following 1 day and 7 days incubation at 37°C. Control: 10% GelMA. Data presented as mean ± standard deviation (SD). *p=0.0136; ***p=0.0009; ****p<0.0001; ns, not significant (*n* ≥ 3). (**d-f**) The influence of gelatin on the stress relaxation properties of PVA hydrogels. Means of normalized stress measurements immediately after casting (**d**) and after 48 h of washing (**e**) are shown. The dashed line indicates the cut-off used for quantifying the stress relaxation time t_0.85_. (**f**) Relaxation times t_0.85_ before (0 h) and after washing (48 h). Data presented as mean ± SD. *p<0.05 (*n* ≥ 2). The 1.5% nPVA group cannot form a gel without gelatin.

Next, we devised a multistep rheological test to mimic the volumetric printing process. As shown in Figure 2b, the resin exhibited a low shear modulus at temperatures above ~25°C (G’ < 2 Pa) but underwent a rapid sol-gel transition upon cooling of the testing system to 20°C. G’ further increased to about 1000 Pa when the temperature was decreased to 4°C (G’’ ~ 10 Pa). At this point, *in situ* UV irradiation was applied to induce the crosslinking via radical-mediated thiol-ene photopolymerization and a rapid increase in shear moduli (G’ > 6000 Pa, G’’ ~ 40 Pa) was observed. Finally, a temperature ramp was used to liquefy the gelatin component by thermal disassembly of the gelatin network, which was confirmed by a slow decrease in both G’ and G’’ when heating the stage to 37°C (Figure 2b, **Figure S1**).

We also compared the physical properties of nPVA hydrogels systematically with a reference hydrogel (10% GelMA) in terms of photo-reactivity, compressive moduli, mass swelling ratio and viscoelasticity as a factor of gelatin release. Photo-rheology results (**Figure S2**) show that the resins are photocrosslinkable in a wide range of nPVA concentrations (1% - 4%). Although 1% nPVA (Mix 1) appeared to be curable, the gels collapsed after 24 h incubation in PBS at 37°C while gels formed from a 1.5% nPVA resin remained stable after incubation. The physical properties of nPVA hydrogels can easily be tuned by adjusting the polymer content from 1.5% to 4% (Figure S2). We observed an increase in both, G’ and G’’, with the increase of nPVA concentration (**Table 1**). By raising the nPVA concentration from 1.5% to 3%, the compressive modulus (*E*) increased from ~6.7 kPa to 27 kPa, whereas the mass swelling ratio significantly decreased from 50 to 35. Compared to the reference, nPVA hydrogels are significantly softer and show a much higher degree of water-uptake (Figure 2c, **Figure S3a**). To study the impact of gelatin release, we measured the compressive moduli of different nPVA hydrogels either directly after UV curing (Figure 2c, 0 h) or following incubation at 37°C for 24 h and 7 days, respectively. The compressive moduli of nPVA hydrogels decreased with increasing incubation time. For instance, the compressive moduli of 1.5% nPVA gels decreased significantly from 6.7 kPa to 2.7 kPa after 7 days incubation at 37°C. We assume that the decrease of compressive modulus is caused by the release of gelatin and network swelling. To test the kinetics of gelatin release, we performed a colorimetric assay of nPVA hydrogels containing fluorescein-labeled gelatin as a reporter. After 24 h incubation in PBS at 37°C, approximately 87% and 84% of gelatin were released from the 1.5% (soft) and 3% (stiff) hydrogels. After 14 days, the percentage increased to 96% and 93%, respectively (Figure S3b). These results confirm that the thermo-sensitive gelatin component can be almost completely removed at 37°C.

We were interested whether the addition of gelatin shows an impact on the viscoelasticity of nPVA hydrogels, and whether this effect would be reversible upon thermal removal of gelatin. Stress relaxation tests under unconfined compression showed that incorporating gelatin confers faster stress relaxation properties immediately after photocrosslinking. The covalently crosslinked 3% nPVA hydrogels without gelatin showed expectedly a mostly elastic behavior and demonstrated a slow stress relaxation time (t_0.85_) of 712.8 ± 188.1 s. Addition of gelatin reduced t_0.85_ to 170.4 ± 121.1 s (Figure 2d). This enhancement in the viscoelastic behavior is lost after incubation in PBS for 48 h at 37°C. The stress relaxation curve of 3% nPVA with gelatin was measured as t_0.85_ = 987.6 ± 293.4 while t_0.85_ of the sample without gelatin remained at 709.3 ± 69.0 s for the pure 3% nPVA gels (Figure 2e-f). This indicates that the gelatin fraction responsible for a temporary increase in stress relaxation properties was removed from the hydrogels during the 48 h incubation in PBS at 37°C. Decreasing the nPVA concentration to 1.5% while keeping the gelatin content constant produced similar initial stress relaxation dynamics and further decreased the initial t_0.85_ to 63.3 ± 15.8 s. Although a decrease in viscoelasticity was observed after 48 h incubation and t_0.85_ increased to 104.7 ± 15.2 s, some stress relaxing properties were retained in this composition which could be especially favorable for cell spreading.

### 2.3. Volumetric Printing of Complex Hydrogel Constructs

After investigating the physical properties of nPVA hydrogels, we set out to fabricate complex permissive hydrogel constructs with varying mechanical stiffness. Using a commercial Tomolite printing system we first optimized the printing parameters using a Dose test based on a 2D projection. This test establishes a relationship between the photocrosslinking efficiency of a given resin and the light source of the printer and is used to determine the threshold laser dose. Below this threshold, the light absorbed by the resin is not sufficient to generate a stable crosslinked network, whereas too high laser doses result in over-exposure (**Figure S4**). The exposure parameters facilitate screening for the “Optimal Dose” (OD, referring to the light dose [mJ cm^−2^] at which the printed structure is closest to the designed structure) for VBP. The OD decreased with the increase of nPVA concentration. These results are consistent with the photo-rheology results discussed before (Figure S2). Notably, at the same LAP concentration the OD value for a 1.5% nPVA resin is obviously lower than that of the reference resin (10% GelMA) (Figure 3d), and an even lower laser dose (~105 mJ cm^−2^) is required for crosslinking a 3% nPVA resin (Figure S4a). The low light dose required for nPVA resins represents a remarkable advantage for bioprinting where over-exposure and excessive reactive species can compromise the cell viability.^[30–31]^

**Figure 3.**
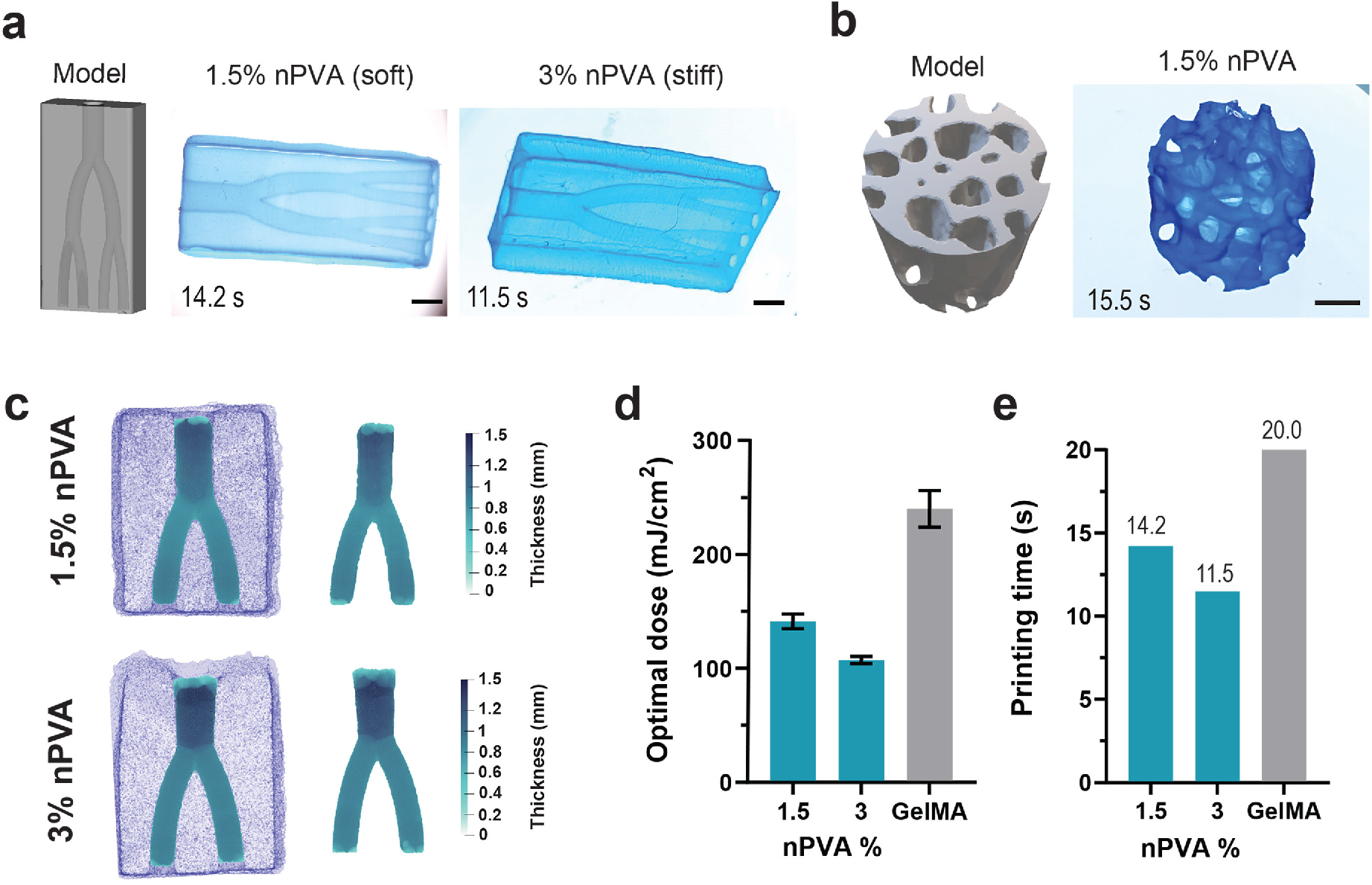
Volumetric printing of perfusable hydrogel constructs. (**a**) CAD model (left) and stereo-microscopic images (right) of perfusable branch constructs fabricated via volumetric printing using a resin containing 1.5% nPVA and 3% nPVA within 14.2 s and 11.5 s, respectively. LAP concentration was 2 mmol m^−3^. (**b**) CAD model (left) and stereo-microscopic image (right) of a trabecular bone femur construct printed with 1.5% nPVA (printing time = 15.5 s). Scale bars, 2 mm. (**c**) 3D volume reconstruction of printed soft and stiff constructs (left) by micro-CT and quantification of the channel diameters (right). Color coding represents the channel diameter. (**d**) The optimal laser dose of different nPVA resins (reference: 10% GelMA) using the branch CAD model. (**e**) The printing time of different nPVA resins and the reference resin using the branch CAD model.

With this information at hand, we further investigated volumetric printing of complex hydrogel constructs. For potential integration with fluid flow through bioreactors, a perfusable branch model was printed with a nPVA concentration as low as 1.5% (**Figure 3a**) within 14.2 s while the 10% GelMA reference resin required a longer time (20 s) to print the same structure (Figure S4c). Increasing the nPVA content further decreased the printing time (Figure 3e). In another example we fabricated a complex trabecular bone construct within 15.5 s using a resin with a polymer concentration of 1.5% nPVA (Figure 3b, **Video S1**). This is an impressive result because fabrication of soft hydrogel constructs with low polymer content leads to softer matrices which are more permissive for 3D cell growth, differentiation, and migration.^[32–35]^ We next employed micro-computed tomography (micro-CT) imaging to characterize the structural fidelity of printed constructs. Figure 3c shows 3D renderings of the volumetrically printed constructs with a bifurcation model. The soft and stiff hydrogel constructs had a channel diameter of 1.34 ± 0.18 mm and 1.45 ± 0.02 mm before the bifurcation, which resulted in two smaller channels of 1.02 ± 0.14 mm and 0.94 ± 0.02 mm respectively (**Figure S5**). These channel diameters are in agreement with the parameters of the 3D stereolithography (.STL) file used for printing. Due to the swelling behavior of printed hydrogel constructs, the width of the printed channels after incubation appears bigger than that of the corresponding CAD model (Figure S5).

### 2.4. Volumetric Bioprinting of Living Hydrogel Constructs

We next evaluated the bio-printability of nPVA resins in terms of cell-compatibility. To shed light on the impact of matrix stiffness on 3D cell growth, two groups of bioresins composed of 1.5% nPVA and 3% nPVA for the soft and stiff group were investigated. Live-dead assays showed that hMSC-laden constructs can be fabricated with high cell viability (~90%) following 2 h, 24 h and 7 days after printing in soft and stiff constructs (**Figure 4a - i, iv**, **Figure S6**). No significant differences were found between the different bioresins (Figure 4b).

**Figure 4.**
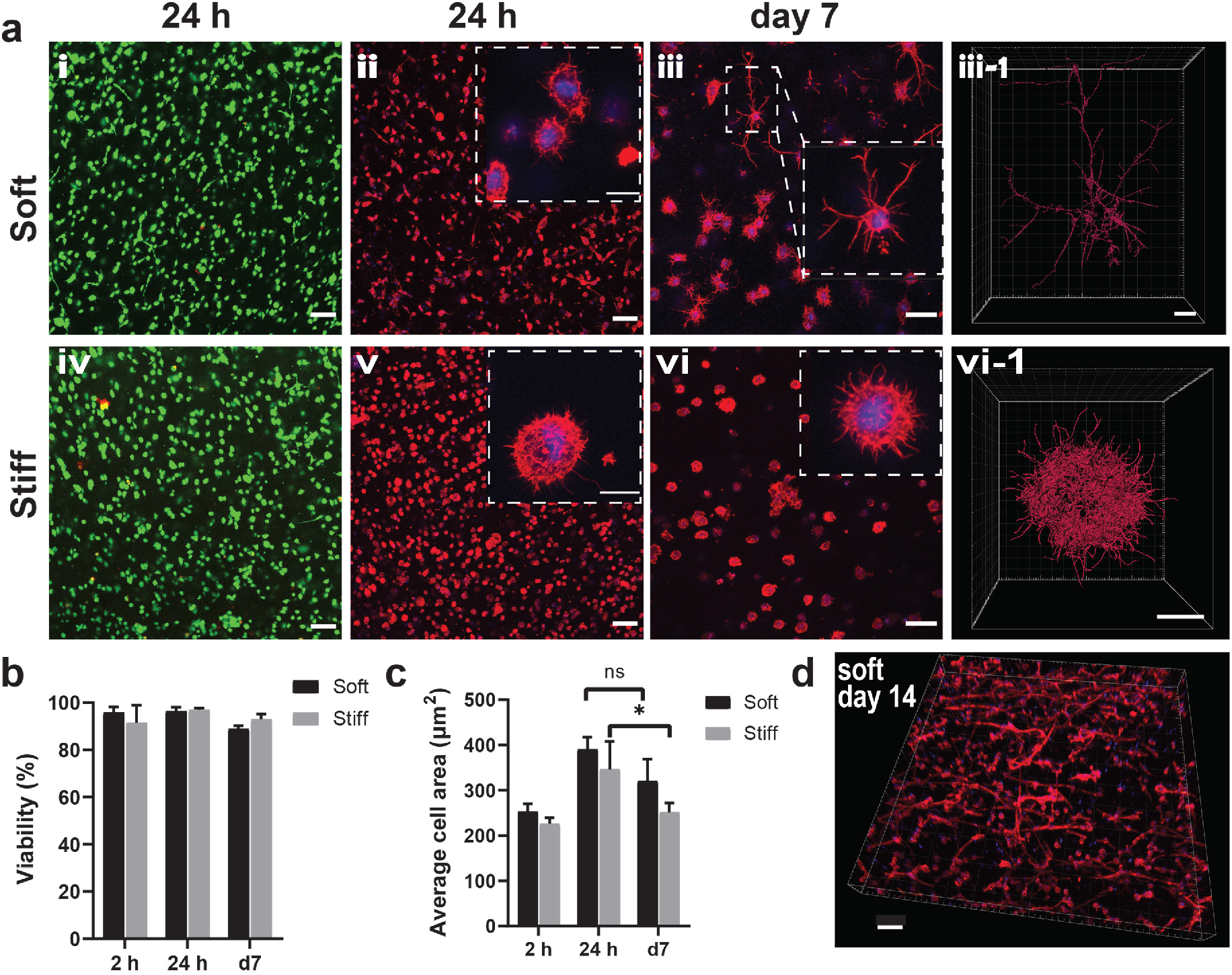
Cell activity in 3D bioprinted dynamic hydrogel environments of varying stiffness. (**a**) Live(green)/dead(red) stained hMSCs following 24 h after printing. Scale bars, 100 μm (i, iv). Confocal images of actin-nuclei stained hMSCs in soft and stiff gels at 24 h (ii,v) and 7 days (iii, vi) after printing. Scale bars, 100 μm (ii, vi) and 50 μm (iii, vii). Scale bars for all inserts are 20 μm. Visualization of single cells in soft and stiff matrix using automated IMARIS dendrite tracking. Scale bars, 10 μm (iii-1, vi-1). (**b**) Quantification of cell viability of hMSCs at different time points. (**c**) Quantification of average cell area at different time points. *p=0.0268; ns, not significant (*n* ≥ 3). (**d**) Confocal image of actin-nuclei stained hMSCs in soft gels following 14 days of osteogenic culture. Scale bar, 100 μm.

Actin-nuclei staining results showed that already 2 h after printing the hMSCs rapidly formed protrusions in the soft matrix, while a much lower degree of spreading was observed in the stiff matrix (**Figure S7**). We reason that the presence of the temporary gelatin network in the matrix and the resulting viscoelasticity (Figure 2d-f) facilitates fast cell spreading and cell-matrix remodeling. After 1 day of osteogenic differentiation, cells in the soft matrix appeared with a more elongated morphology and even displayed cell-cell contacts, contrary to cells in the stiff matrix (Figure 4a - ii, v). After 7 days of cultivation, more interconnected cells with long protrusions were observed in the soft matrix, while more single cells with short protrusions were found in the stiff matrix (Figure 4a - iii,vi, **Video S2**). Automated dendrite tracking was used to compare the morphological differences of single cells in the soft and stiff matrices. A stellate cell morphology was observed within the soft matrix, whereas cells embedded in the stiff matrix exhibited much shorter protrusions. Over time, cells continued 3D morphogenesis and an interconnected network was observed in the soft matrix after 14 days (Figure 4d).

Interestingly, the average cell area initially increased between 2 h and 24 h after printing, but then decreased after 7 days (Figure 4c). This observation implies remarkable morphological changes of bone cells during 3D osteogenic differentiation considering mature bone cells, termed osteocytes, typically have an ellipsoid cell body and long dendrites in native tissues. The transition from an osteoblastic to an early osteocytic phenotype is associated with the reduction of cell area.^[36]^ Alizarin red staining evidenced mineral deposition within the soft and stiff matrices following 21 days of *in vitro* osteogenic culture (**Figure S8a**), indicating the progress of matrix mineralization as a hallmark of osteogenic differentiation. No statistical difference between soft and stiff matrices was observed (Figure S8b). Nevertheless, further work is necessary to probe the cellular and molecular dynamics in these printed environments.

To make full use of the permissive properties of the soft bioresin we tested the fabrication of more complex constructs. As shown in **Figure 5**, a permissive hMSC-laden construct was successfully printed using a branch model within 15 s. Interestingly, cells in the printed branch model could self-organize into irregularly shaped multicellular aggregates and migrate towards the hollow space of the inner channels after printing (**Figure S9**). Live-cell confocal images revealed that hMSCs formed smaller aggregates in the smaller channels (C2 and C3) while bigger cell aggregates with round shapes were observed in the larger C1 channel (Figure 5b-d).

**Figure 5.**
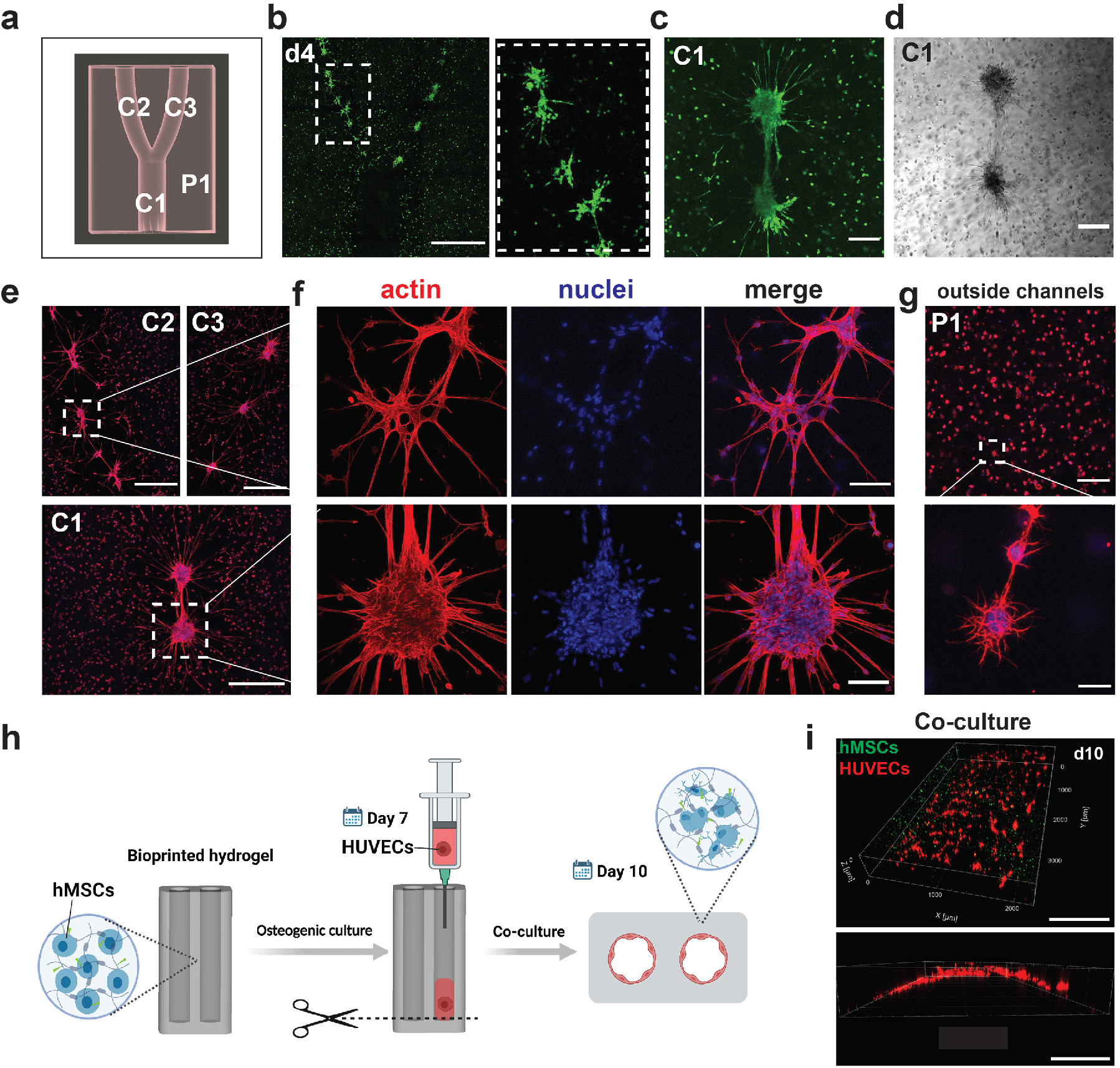
VBP of perfusable cell-laden constructs. (**a**) The simplified CAD model containing a tubular structure was used for VBP of perfusable constructs. The larger channel C1 splits into the smaller channels C2 and C3, while the surrounding area is referred to P1. (**b**) Confocal images of Calcein AM-stained cells (green) at day 4 in a construct printed from the soft bioresin. Scale bar, 1 mm. (**c-d**) Confocal and bright-field images of Calcein AM-stained cell aggregates adjacent to the big channel (C1) on day 4. Scale bars, 200 μm. (**e**) Confocal images of actin-nuclei stained cell aggregates at positions C2, C3 and C1 on day 7. Scale bars, 500 μm. (**f**) High magnification confocal images of two actin-nuclei stained cell aggregates in the small channels (C2, top) and big channel (C1, bottom). Scale bars, 100 μm. (**g**) Confocal actin-nuclei images of representative cells outside the channels (P1) in the printed construct. Scale bars, 100 μm (top) and 20 μm (bottom). (**h-i**) VBP of a pre-vascularized model. (**h**) Illustration of the experimental steps: Using a custom simplified CAD model, a hMSC-laden construct is bioprinted and differentiated under osteogenic conditions until day 7. Then, HUVECs in a supporting collagen matrix are injected into the channels. Over time, the collagen hydrogel is remodeled and HUVECs can self-organize into a cell layer in the channels of hMSC-laden constructs after another 3 days. (**i**) Confocal images of hMSCs and HUVECs stained with calcein-AM and DiD, respectively, after co-culture for 3 days. Scale bars, 1 mm.

Actin-nuclei staining was used to detail the location and morphology of cell aggregates after 7 days of culture. The cell aggregates aligned with the orientation of the channels. Besides, the spreading of actin-based cell protrusions into all directions from the hMSC aggregates was observed (Figure 5e-f, **Figure S10**). No aggregates were found for cells outside the channels (P1) even though they formed local cell-cell contacts (Figure 5g). We reason that cells surrounding the channels have more access to nutrients, as well as more physical freedom to self-organize into cell aggregates. In addition, these results exemplify that geometrical cues play an important role in generating aligned multicellular systems.^[37–38]^ Therefore, our nPVA bioresin in combination with VBP may find future applications in complex tissue engineering. In biofabrication, 3D construction of hetero-cellular environments will provide the means to better mimic native tissue environments where paracrine signaling is crucial for cell-cell communication and tissue development.^[39–40]^ To test if the nPVA bioresins can be used for 3D co-culture applications, we seeded human umbilical vein endothelial cells (HUVECs) into the hollow channels of a 3D printed hMSC-laden construct (printing time = 12 s) to generate a pre-vascularized model. Before co-culture, the hMSC-laden hydrogel construct was osteogenically differentiated for 7 days. After being dispersed in a supportive collagen hydrogel, HUVECs were loaded into the channels, followed by 3D co-culturing for another 3 days (Figure 5h). Confocal images evidenced the formation of an endothelial layer (Figure 5i).

### 2.5. Volumetric 4D Photopatterning

Finally, our interest focused on fast tomographic volumetric photopatterning and if it can be used to immobilize molecules-of-interest within a preformed hydrogel for potential spatiotemporal control of cell-matrix interactions during 3D cell culture. We expected that non-reacted norbornene groups in a nPVA hydrogel formed by an off-stoichiometric crosslinking are accessible for radical-mediated thiol-ene photo-conjugation^[41–44]^ using tomographic photopatterning. This 3D patterning process is realized in three steps as shown in **Figure 6**. First, a centimeter-scale hydrogel was generated by tomographic volumetric printing with a 3% nPVA resin (Mix 4). Subsequently, the construct was immersed in a reaction buffer containing thiolated fluorescein (FITC-SH, 0.85 mmol m^−3^) and LAP photoinitiator (1.7 mmol m^−3^). The FITC-SH molecule was immobilized within predetermined 3D regions by performing a second volumetric printing procedure. The printed 3D patterns were visualized using a fluorescent stereomicroscope after the removal of soluble dye molecules through washing.

**Figure 6.**
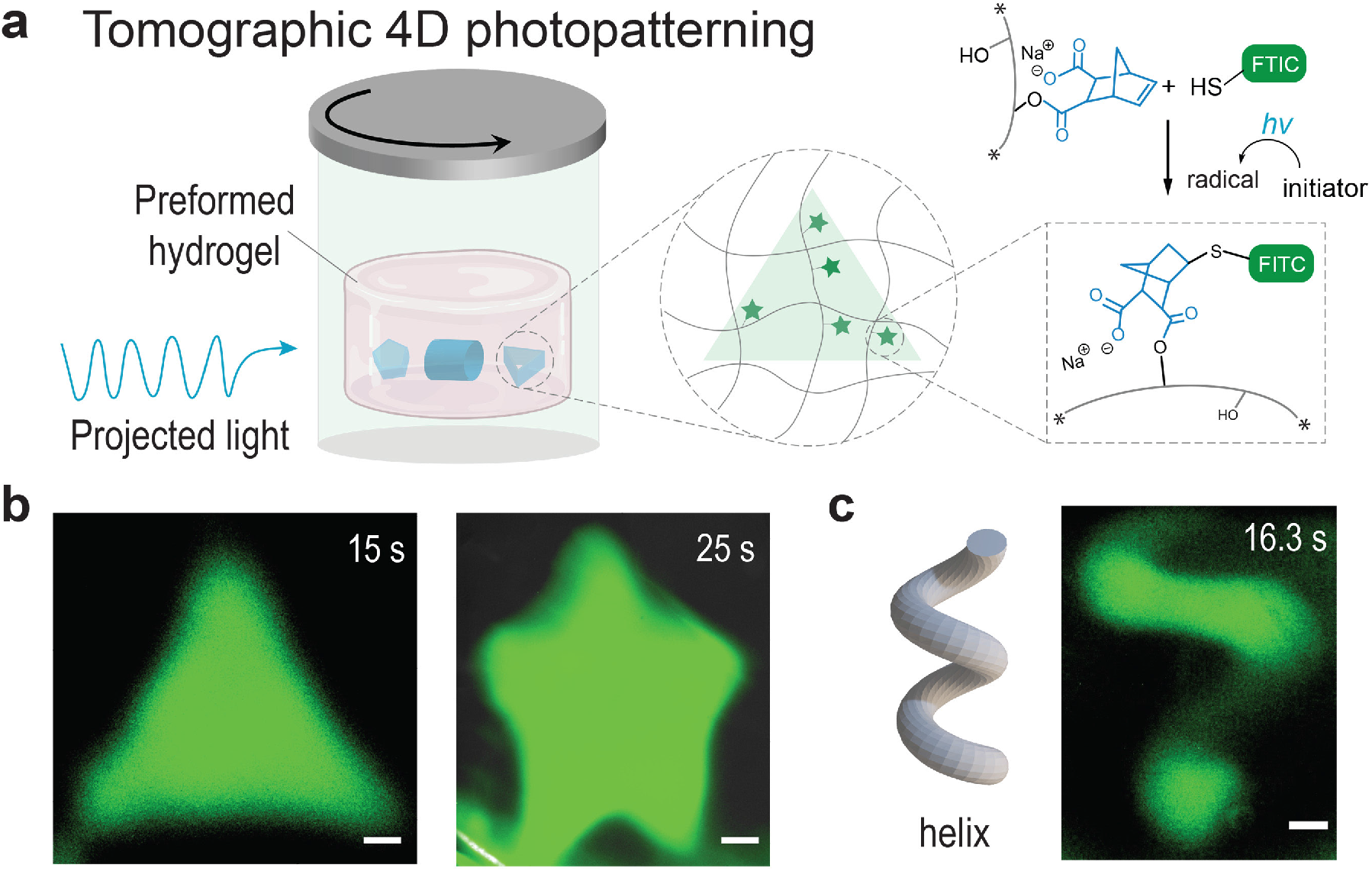
Tomographic volumetric photopatterning of chemical cues in preformed hydrogel constructs within seconds. (**a**) Illustration of volumetric thiol-ene photopatterning process. (**b**) Top view of fluorescent stereomicroscopy images of a patterned triangular prism and five-star models. (**c**) A complex 3D helix model was patterned within 16.3 s: CAD model (left) and stereomicroscopic image of the patterned construct (right). Scale bars, 500 μm.

A 3D triangular prism was created inside the printed construct through tomographic thiol-ene photopatterning within 15 s. In addition, geometrically complex 3D patterns such as star-shaped cylinder (Figure 6b) and a free-form helix (Figure 6c, **Video S3**)^[45]^ were printed within 25 s and 16.3 s, respectively. Of note, a printed cylinder with a line channel could be site-selectively functionalized with the FITC-SH molecule around its channels within 18 s (**Figure S11**). Given the short patterning time and the cell-friendly light wavelength (405 nm), we are convinced that the described volumetric 4D photopatterning will open up new possibilities in fabricating biomimetic 3D tissue models. Future work can harness this methodology to functionalize distinct parts of the printed matrices to guide cellular behavior and tissue morphogenesis in a spatially and temporally controlled fashion.

## 3. Conclusion

We developed a synthetic dynamic photoresin based on nPVA for VBP of functional ultrasoft hydrogel constructs within 7-15 seconds, thereby circumventing the limitations of existing resin materials. This system utilizes highly efficient thiol-norbornene photo-click chemistry to generate modular hydrogel environments with nPVA concentrations as low as 1.5%. Thereby the addition of a temporary gelatin network improves the printability of the resin and enables us to decouple the physicochemical properties of printed matrices from the fabrication process. Using this resin, we successfully fabricated stable stress-relaxing hydrogel constructs with low polymer concentration that support fast cell spreading and 3D cell growth, osteogenic differentiation of stem cells and matrix mineralization, the formation of aligned multicellular aggregates and co-culture with endothelial cells. We report the fast site-specific grafting of chemical cues within preformed hydrogels by tomographic thiol-ene photopatterning. These features are up to now unachieved in conventional gelatin-based resins. In the future nPVA resins may enable high-fidelity VBP of permissive tissue constructs using a recently reported optical-tuning technique to counteract light scattering.^[46]^ We are convinced that the reported synthetic dynamic photoresin will path the way to new applications for fast biomanufacturing of tailored 3D tissue and organ models for regenerative medicine.

## 4. Experimental Section

### Synthesis of nPVA

nPVA was synthesized as previously described.^[47]^

### Volumetric Printing Procedure

Resins were formulated as described in Supplementary Table 1. Briefly, 0.8-1.2 mL of resin was transferred into the printing glass vials (diameter: 10 mm). The vials were chilled at 4°C for 5-10 min to form gels by physical crosslinking of sacrificial gelatin component. Then, printing was carried out on a commercially available Tomolite v1.0 volumetric printer (Readily3D, EPFL Innovation Park, Switzerland) equipped with the latest version of Apparite software.^[7]^ After printing, the glass vials were warmed to 37°C to remove the unpolymerized resin. Thereafter, the printed object was washed with prewarmed PBS. Printed objects were transferred into a solution containing sodium persulfate (SPS, 5 × 10^−3^ mol m^−3^) and tris(2,2-bipyridyl) dichlororuthenium(II) hexahydrate (Ru, 0.5 × 10^−3^ mol m^−3^) for post-curing under a 405 nm LED lamp (Thorlabs, Germany, 10 mW cm^−2^) for 5 min. The printed branch constructs were perfused with Alcian blue to facilitate visualization. Constructs were imaged on a Leica M205 FA stereomicroscope coupled with a DFC550 CCD camera for real color imaging.

### Micro-computed tomography (micro-CT)

The 3D renderings were obtained with micro-CT scans performed in air with a micro-CT45 device (Scanco Medical, Bassersdorf, Switzerland). Before scanning the water was carefully removed from the channels using tissue papers. The printed constructs were scanned at 45 kVp energy and 177 μA current, using an integration time of 600 ms. The resulting 3D images had a voxel size of 17.2 μm. A Gaussian filter (σ = 1.2, support 1) was used to reduce noise in the images before converting the grayscale images to binary images using a threshold corresponding to −10 mg hydroxyapatite cm^−3^. A negative threshold was needed because the scan was made in air where the minimum value corresponds to ca. −167 mg HA cm^−3^. The greatest connected component was chosen to remove structures not belonging to the printed construct. The low attenuation property of the hydrogel required additional three binary erosion and dilation iterations to smooth the surface. The channel thickness was analyzed by generating a rectangular volume of interest, laying completely within the boundaries of the 3D image, and the distance transformation method.^[48]^ The image processing was performed using an in-house developed software framework based on Python 3.6.6 (Python Software Foundation) using the SciPy 1.2.0 library and IPL software (V5.42 Scanco Medical). The 3D renderings were generated with ParaView (V5.10.1, Kitware Inc.) and channel widths were evaluated by using ParaView’s built-in measurement tools.

### Volumetric Bioprinting (VBP) and Cell Culture

The nPVA-gelatin bioresins were prepared as indicated above except the addition of cells and 2 mmol m^−3^ CGRGDSP peptide. All components were sterile-filtered (0.2 μm) except the commercially available sterile gelatin (Sigma-Aldrich, cat. no. G2500). Human bone marrow-derived mesenchymal stem cells (hMSC) (pooled donor, Lonza, Switzerland) were harvested at passage 3-4 and resuspended in 0.8 mL of bioresin at a concentration of 5 × 10^6^ cells mL^−1^. The well-mixed bioresin was transferred into autoclaved glass vials and immediately chilled at 4°C. After printing, the non-crosslinked resin was liquefied at 37°C and the cell-laden hydrogel constructs were retrieved and washed with prewarmed PBS under sterile conditions, followed by cutting the printed constructs into 12 pieces (radius: 3 mm; thickness: 2 mm) for further evaluation. Cell viability was monitored at different time points following cultivation in osteogenic medium composed of Dulbecco’s Modified Eagle’s Medium (DMEM), 10% fetal bovine serum (FBS), 1% antibiotic-antimycotic solution, ascorbic acid (50 μg mL^−1^), dexamethasone (100 nmol m^−3^) and beta-glycerophosphate (10 mmol m^−3^). Medium was changed every 2 days.

### Perfusable Cell-laden Constructs and Co-culture

Perfusable cell-laden constructs were fabricated using the branch CAD model following the standard VBP procedure with a seeding density of 2.5 × 10^6^ hMSCs mL^−1^. The printed constructs were stained by incubating in Calcein-AM (2 μmol m^−3^) at 37°C and 5% CO_2_ for 2 h. Live-cell imaging was performed using a confocal microscope (Leica SP8) after cultivation for 3-4 days.

The hMSC-laden models with hollow channels (diameter = 1 mm) were cultivated in osteogenic medium for 7 days. Constructs were stained in Calcein-AM (2 μmol m^−3^) for 1.5 h at 37°C and 5% CO2 before seeded with human umbilical vein endothelial cells (HUVECs, pooled donor, Lonza, Switzerland). Endothelial Cell Growth Medium-2 (EGM-2, Lonza, Switzerland) was used as the medium. The harvested HUVECs (passage 3) were labelled with DiD cell labelling solution (Vybrant, Thermo Fisher Scientific), and seeded with a blunt-tip cannula (diameter = 0.8 mm) into the perfusable channels at a density of 5 × 10^6^ cells mL^−1^ with a supporting collagen type I matrix (Corning, 1 mg mL^−1^). The constructs were cultured in a co-culture medium (1:1 mix of osteogenic medium and EGM-2).

### Volumetric 4D Photopatterning

Cylindrical hydrogel constructs were printed using the 3% nPVA resin and washed for 3 days at 37°C. Then the printed cylinders were immersed in a solution containing 0.85 mmol m^−3^ FITC-SH and 1.7 mmol m^−3^ LAP overnight at 4°C. To improve the stability of thiolated molecules, TCEP (21 mmol m^−3^) and pyrogallol (85 ppm) were added to the working solution. The stained hydrogel constructs were transferred to glass vials to perform volumetric thiol-ene photopatterning by conducting a second printing step, followed by washing in PBS on a shaker for two days to remove non-reacted FITC-SH in the printed constructs. The patterned objects were imaged with a Leica M205 FA stereomicroscope equipped with an X-Cite 200DC lamp with 3 mm light guide and widefield epifluorescence filter sets.

## Supporting Information

Supporting Information is available from the author.

## Acknowledgements

This project was partially supported by the Swiss National Science Foundation (SNSF) Project funding (No. 188522). X.H. Q. acknowledges the financial support by the SNSF Spark Award (No. 190345). W. Q. gratefully acknowledges financial support from China Scholarship Council (No. 202006790027). The authors would like to thank Drs. Paul Delrot and Damien Loterie (Readily3D) for their generous technical support as well as the staff members of ScopeM at ETH Zurich for their assistance with imaging. Dr. Riccardo Levato (Utrecht University), Dr. Xiaopu Wang and Prof. Bradley Nelson (ETH Zurich) are acknowledged for providing the STL files.

## Notes

### Competing Interest Statement

The authors have declared no competing interest.

### Summary of Updates

- title - Abstract

