## Supporting Information for "A Synthetic Dynamic Polyvinyl Alcohol Photoresin for Fast Volumetric Bioprinting of Functional Ultrasoft Hydrogel Constructs"

**This PDF file includes:**

Supplementary Methods

Supplementary Figures 1 to 12

Supplementary Table 1

Supplementary References

**Other Supplementary Materials for this manuscript include the following:**

Videos S1 to S3

**Supplementary Methods**

*Materials and reagents.* All chemicals were purchased from Sigma-Aldrich unless otherwise indicated. All reagents were used as received unless otherwise noted.

*Synthesis of Norbornene-functionalized Polyvinyl Alcohol (nPVA).* Norbornene-functionalized polyvinyl alcohol (nPVA) was prepared as previously reported.[1] PVA (8 g, 47 kDa, cat. no. 10853) was dissolved in anhydrous DMSO (130 mL) under argon protection at 60°C, and then 10% of solvent was distilled under a fine vacuum. Then, p-toluenesulfonic acid (pTsOH, 18.2 mg) was dissolved in anhydrous DMSO (0.5 mL) and transferred into PVA solution under argon atmosphere. Cis-5-Norbornene-endo-2,3-dicarboxylic anhydride (3 g, abcr GmbH) was dissolved in anhydrous DMSO (15 mL) and was added into the PVA/pTsOH solution dropwise. After a 24 h reaction at 55°C, the solution was cooled down to room temperature and transferred into a dialysis tube ($M_w$ cut-off, 6-8 kDa). After dialysis against $NaHCO_3$ (pH 8.0) for 24 h and then against MilliQ water for 24 h, the product was lyophilized to yield 9.0 g of nPVA. The degree of substitution was 7.8% as determined by $^1$H-NMR in $D_2O$ (Figure S12).

*Synthesis of Gelatin Methacryloyl (GelMA).* GelMA was synthesized by reacting gelatin (Type A, porcine skin, cat. no. G2500) with methacrylic anhydride (cat. no. 276685) as previously described.[2] The degree of substitution (DS) of GelMA was estimated with $^1$H-NMR (Bruker Ultrashield, 400 MHz) in $D_2O$. Integration signal (2.87–3.00 ppm) in GelMA was compared to unmodified gelatin lysine integration signal (2.87–3.00 ppm). Phenylalanine signal (7.1–7.4 ppm) was used as internal reference. DS was found to be ≈57% using the following equation 1:

DS = [1- (lysine methylene proton of GelMA)/(lysine methylene proton of gelatin)]*100% (1)

*Hydrogel Formation and Characterization.* The nPVA-gelatin hydrogel formulations were composed of gelatin, photocrosslinkable components nPVA and 2 kDa poly(ethylene glycol) dithiol PEG2SH (Laysan Bio Inc. USA), photoinitiator lithium phenyl-2,4,6-trimethylbenzoylphosphinate (LAP) and pyrogallol. In all tested groups, the concentration of gelatin, LAP and pyrogallol were 5% w/v, 2 mmol $m^{-3}$and 50 ppm, respectively. Designed testing groups were based on the constant stoichiometric ratio (4:5) between thiol and norbornene groups for varying nPVA concentrations (1%, 1.5%, 2%, 3%). The formulations are listed in Table S1. For the reference resin, GelMA was dissolved in 2 mM LAP solution at 37°C to form a 10% solution.

*Photo-rheology Measurement.* Oscillatory sweeps were carried out on a modular rheometer MCR 302 (Anton Paar, Graz, Austria) with a 20 mm parallel plate to determine the thermal responsivity of nPVA resins with and without 5% gelatin. The complex modulus ($G^*$) was determined using equation 2:

$$G^{*2} = G'^2 + G''^2 \tag{2}$$

The temperature was set for a ramp change between 37°C and 4°C at a rate of 1°C min$^{-1}$. To mimic the bioprinting process, the temperature-controlled rheometer was equipped with a 365 nm UV-LED lamp (Thorlabs, Germany) for *in situ* photopolymerization. The temperature was initially set at 37°C and then decreased to 4°C at a rate of 1°C min$^{-1}$; next, 365 nm UV-LED light was introduced at 10 mW cm$^{-2}$; and the sample was heated back to 37°C at a rate of 1°C min$^{-1}$. All measurements were performed in triplicates (n=3) at 1% stain and 1 Hz frequency with a gap thickness of 100 μm. Mineral oil was added around the parallel plate to prevent sample from drying during the testing.

*Mechanical Testing.* The unconfined compression moduli of casted hydrogel disks (diameter, 6 mm; height, 1.8 mm) was measured on a Zwick material testing machine (Zwick 1456, Ulm, Germany) with 10 N load cells at room temperature. The samples were preloaded at 5 mN, then subjected to compressive strain at a rate of 0.01 mm s$^{-1}$. The compressive modulus was determined by the slope in the linear region of the stress-strain curve, which was between 5% and 15% deformation for all samples. A sample size of 3-4 was used.

*Stress Relaxation Testing.* To determine the stress relaxation kinetics, unconfined compression tests were performed on a Texture Analyser TA.XTplus (Stable Micro System, USA). Casted hydrogel disks of 6 mm diameter and 1.8 mm height were compressed at a test speed of 0.02 mm s$^{-1}$ up to a strain of 15%. Then the strain was held constant and the stress was recorded over a time period of 20 min. Stress relaxation time $t_{0.85}$ was defined as the time needed to reach 85% of the initial stress measured and was chosen for quantitative comparisons to include all tested samples. Measurements were repeated after washing the gels in PBS for 48 h at 37°C to remove the gelatin fraction. Multiple paired t tests with Holm-Šídák correction were performed to compare the two time points for each gel composition.

*Synthesis of FITC-gelatin.* 200 mg gelatin (Type A, porcine skin, cat. no. G2500) was dissolved in 10 mL phosphate buffered saline (PBS) at 40°C. NHS-fluorescein (6.5 mg) was dissolved in 1 mL dimethyl sulfoxide (DMSO), and then added into the gelatin solution and mixed thoroughly by magnetic stirring. The mixture was stirred at room temperature for 24 h under light protection. The yellow solid material (~180 mg) was obtained after dialysis against MilliQ water for one week and lyophilization.

*Gelatin Release.* A hydrogel cylinder model (diameter, 6 mm; height, 1.8 mm) was used to test the release of gelatin fraction from photocrosslinked nPVA-gelatin hydrogels. Typically, 60 µL of nPVA-gelatin-fluorescein (gelatin-fluorescein: gelatin = 1:14) resins were casted in a multiwell PDMS mold by photocrossinking for 5 min (365 nm, 15 mW $cm^{-2}$). The hydrogels were transferred to a 24 well plate filled with 1 mL of PBS for incubation at 37°C, wrapped with parafilm. At given time intervals, 500 µL of supernatants were collected and replaced with 500 µL of fresh PBS. After the final collection, the remaining solution together with the hydrogel cylinder was homogenized by using pellet pestles for 10 min. The fluorescence intensities of all the harvested solutions were measured at 535 nm (excitation, 485 nm) using a plate reader (Tecan M200, Zurich, Switzerland). The percentage of release at each time point was determined by normalization to the cumulative amount released from the samples.

*Live/dead Staining.* Fluorescent live/dead assays were performed to determine cell viability in 3D printed constructs (2 h, 24 h and 7 days after printing; n=4). Constructs were washed twice with PBS and stained for 25 min at 37°C and 5% $CO_2$ with a solution containing Calcein-AM (2 µmol $m^{-3}$) and ethidiumhomodimer-1 (EthD-1) (2 µmol $m^{-3}$) in PBS for live cells (green) and dead cells (red), respectively. A confocal laser scanning microscope (Leica SP8, Germany) was used for image acquisition and images were analyzed using the automated particle analysis plugin on ImageJ (NIH).

*Actin-nuclei Staining.* Cell morphology was evaluated at different time points after fixation with 4% paraformaldehyde (PFA) for 20 min at room temperature. Unspecific binding sites were blocked with 1% bovine serum albumin (BSA) for 2 h in PBS and cell membranes were permeabilized with 0.2% Triton-X100 in PBS for 10 min. Samples were washed with 0.1 % BSA in PBS for 3 times. Cell nuclei and the actin cytoskeleton were stained with Hoechst 33342 (1 µg $mL^{-1}$) and Phalloidin-TRITC (2 µmol $m^{-3}$) in PBS supplemented with 0.1% BSA for 2 h at room temperature. Samples were washed by PBS and imaged on a Leica SP8 confocal microscope equipped with 10× and 25× objectives. Quantification of cell area was performed in ImageJ by evaluating the actin area and nuclei number.

*Alizarin Red Staining (ARS).* Alizarin Red S dissolved in distilled water (2 mg $mL^{-1}$, pH 4.22) was applied to stain the mineral deposition in osteogenically cultured samples for 30 min at room temperature. Then, samples were washed 5 times with distilled water with 5 min between each washing step and imaged on a Leica DMi1 optical microscope. Quantification of ARS was performed with images taken at 10× magnification for each group using ImageJ. A color threshold was applied by selecting only the red channel in RGD color space and the intensity of red color was measured (n=3).

*Statistical Analysis.* Results were reported as mean ± standard deviation (SD). Statistical analyses were conducted using GraphPad Prism 9.2.0. Student's t-test was applied to analyze the statistical significance between two groups. One-way ANOVA was used to analyze the statistical significance between three groups.

**Figure S1**

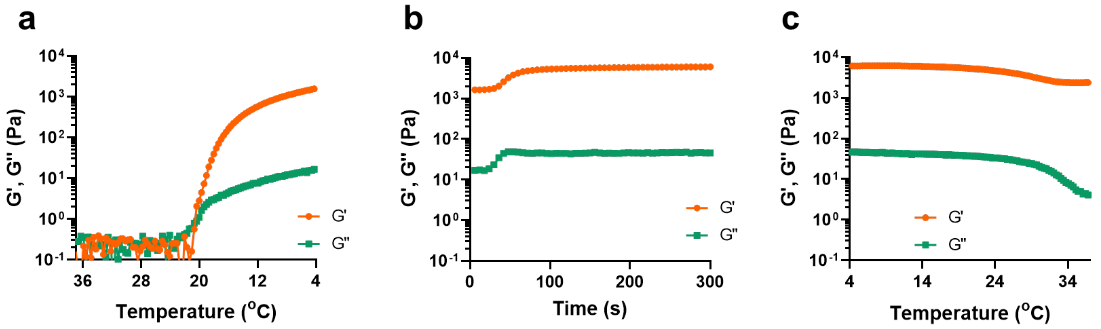


**Figure S1.** Multi-step rheology test of the nPVA photoresins. Storage (G') and loss (G'') moduli in a multi-step rheology measurement of 2% nPVA mimicking the volumetric printing process: (**a**) Stage 1: the sample was cooled from 37°C to 4°C at a rate of 1°C $min^{-1}$ to reach a solid state; (**b**) Stage 2: the sample was photocrosslinked under UV exposure for 5 min (365 nm, 10 mW $cm^{-2}$) at 4°C; (**c**) Stage 3: the crosslinked sample was heated back from 4°C to 37°C at a rate of 1°C $min^{-1}$. Strain, 1%; frequency, 1 Hz. PEG2SH (2000 Da) was used as the crosslinker for a final thiol:ene molar ratio of 4:5. The concentration of photoinitiator LAP was 2 mmol $m^{-3}$.

**Figure S2**

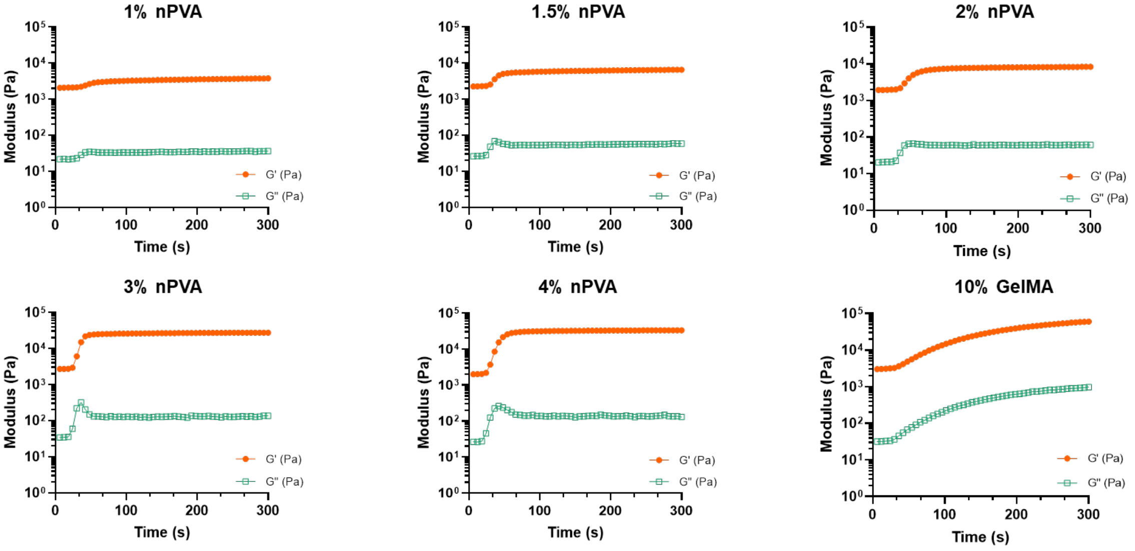


**Figure S2.** Photo-rheology test of resin formulations with varying polymer concentration. Storage (G') and loss (G'') moduli during an oscillatory time sweep of different resins. The sample was pre-cooled at 4°C and subsequently exposed to UV light (365 nm, 10 mW cm$^{-2}$) for 5 min. Strain, 1%; frequency, 1 Hz. The nPVA resins contain 2 mmol m$^{-3}$ LAP, 5% sacrificial gelatin and a stoichiometric amount of PEG2SH crosslinker (thiol:ene = 4:5), whereas the reference resin contains 10% GelMA and 2 mmol m$^{-3}$ LAP.

**Figure S3**

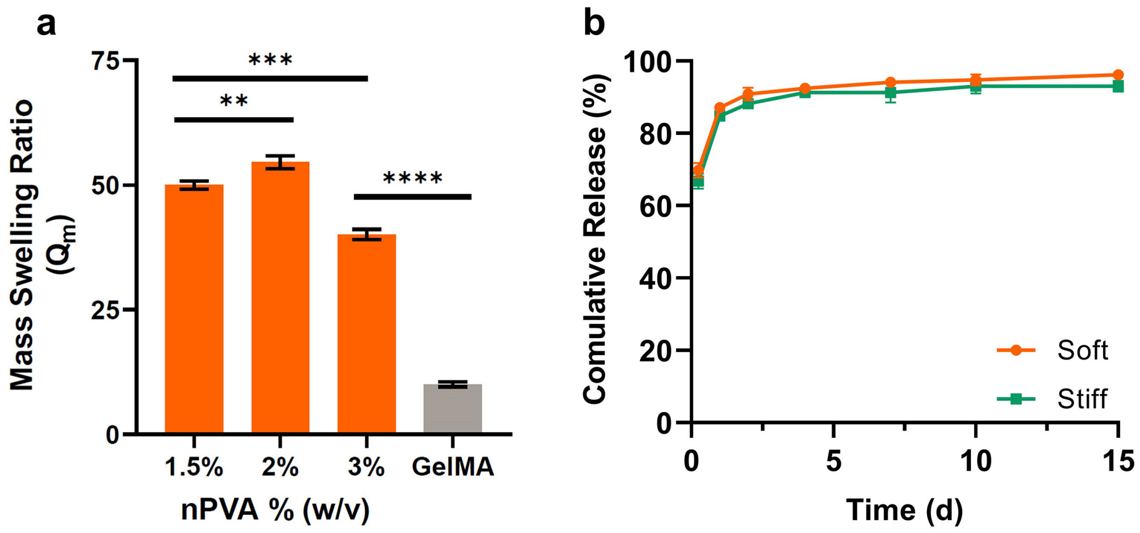


**Figure S3.** Mass swelling ratio of nPVA hydrogels and time-resolved release of sacrificial gelatin. (**a**) Equilibrium mass swelling ratio ($Q_m$) of different nPVA hydrogels versus GelMA hydrogels. **p=0.0065; ***p=0.0002; ****p<0.0001 (n ≥ 3). (**b**) Accumulated release of sacrificial gelatin from 1.5% nPVA (soft) and 3% nPVA (stiff) hydrogels after incubation at 37°C at different time points for up to 15 days.

### Figure S4

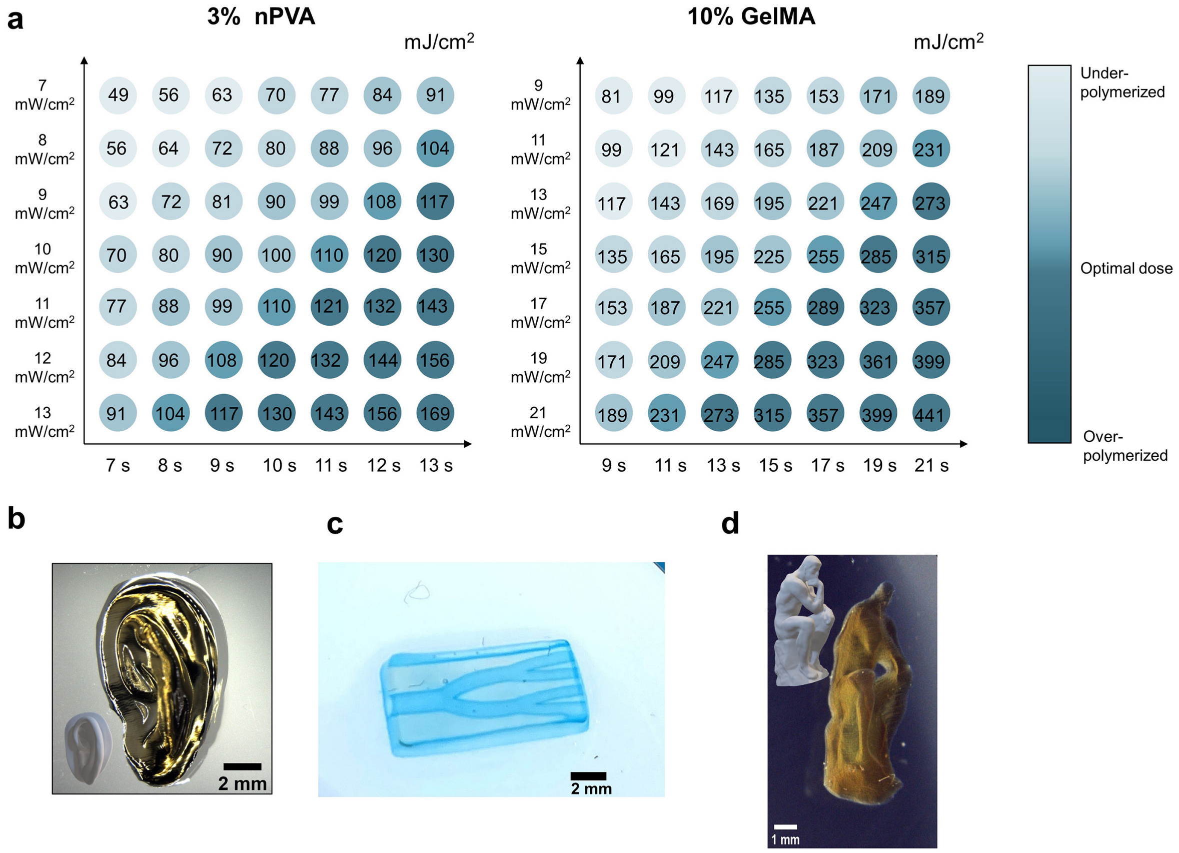


**Figure S4.** Printability characterization. (**a**) Dose test diagrams of 3% nPVA and 10% GelMA resins showing their printability in function of laser dose. Color code indicates the printability of the resin for volumetric printing. (**b**) Microscopic image of an anatomic ear construct printed with 3% nPVA (printing time = 10.3 s). (**c-d**) Microscopic images of a perfusable branch construct (c) and a Thinker construct (d) using 10% GelMA with a printing time of 20.0 s and 33.1 s, respectively.

Figure S5

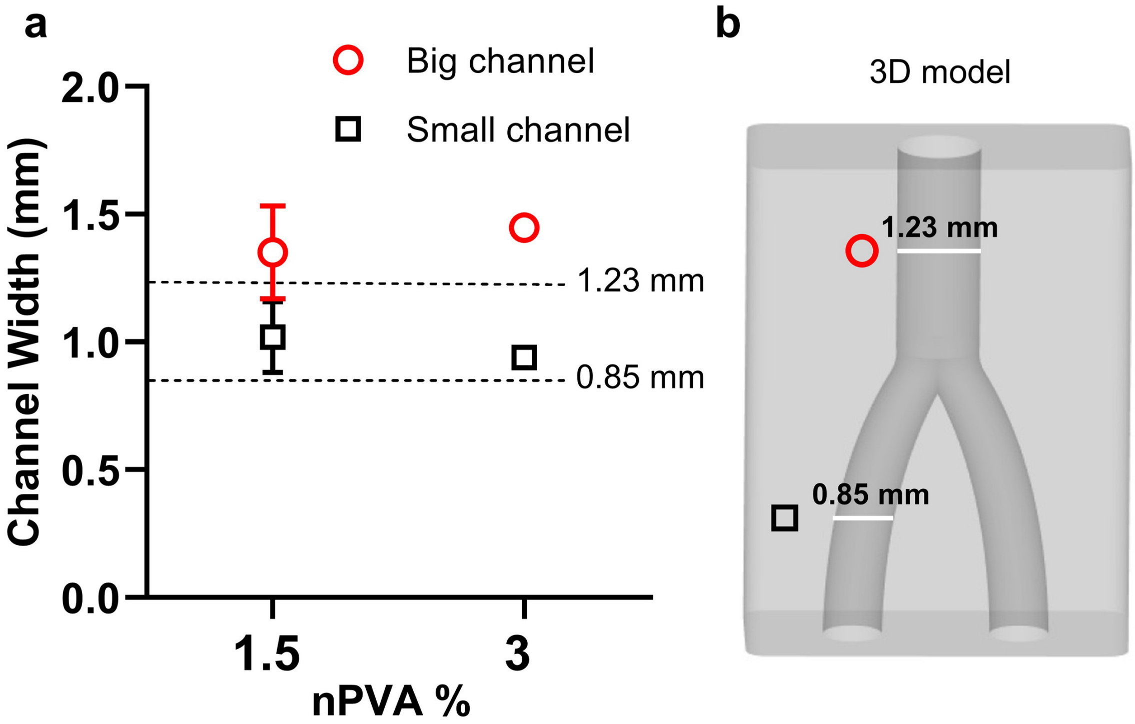


**Figure S5.** Quantification of structure fidelity in printed constructs with micro-CT. (**a**) Quantification of the channel width of the printed branch constructs at different positions. (**b**) The CAD model used for printing.

**Figure S6**

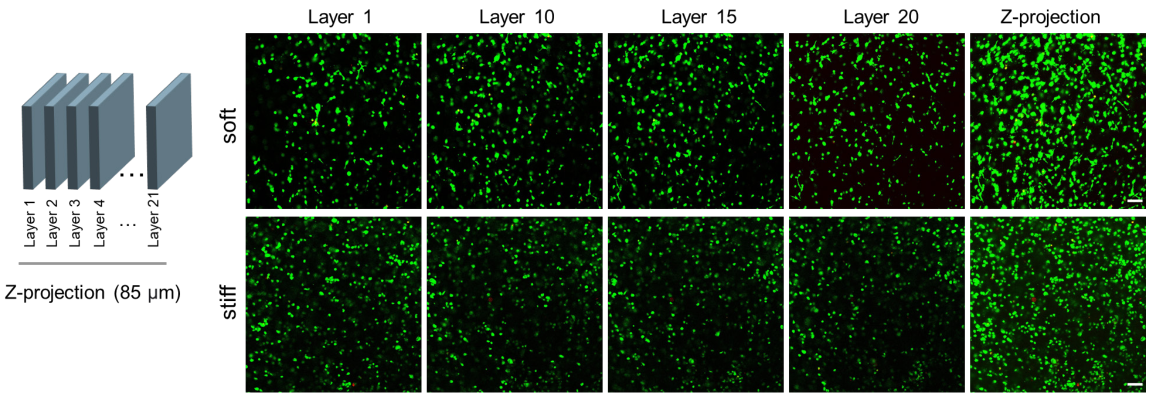


**Figure S6.** Cellular spatial distribution in the bioprinted constructs. Confocal images of hMSCs distributed in soft and stiff nPVA hydrogels after VBP at different depths. Green, calcein-AM; red, ethidium homodimer-1. Scale bars, 100 μm.

**Figure S7**

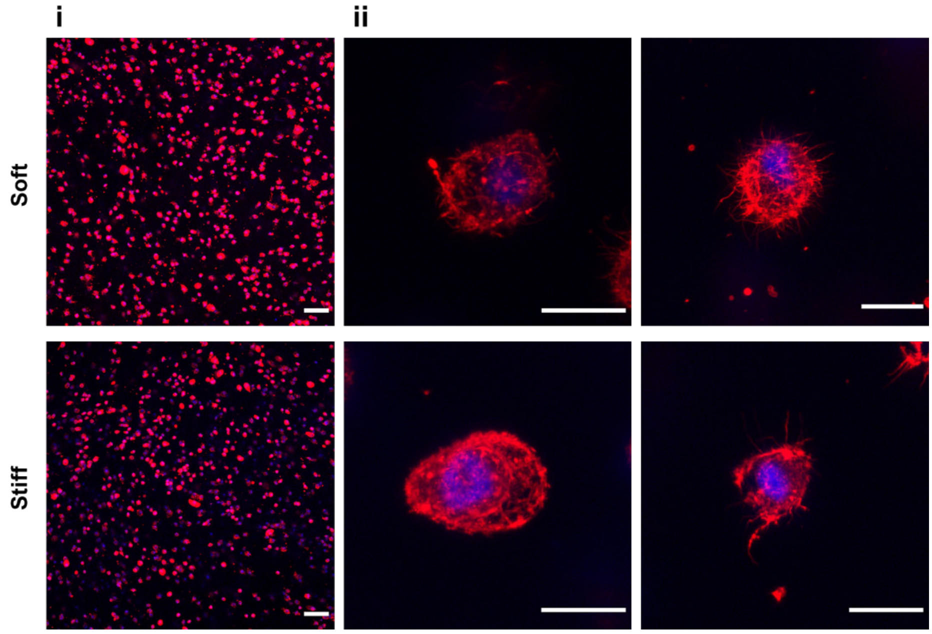


**Figure S7.** Cell morphologies after VBP. Confocal microscopy images of representative hMSCs in soft and stiff hydrogels stained with nuclei (blue) and actin (red) following 2 h after VBP. Scale bars, 100 μm (i) and 20 μm (ii).

**Figure S8**

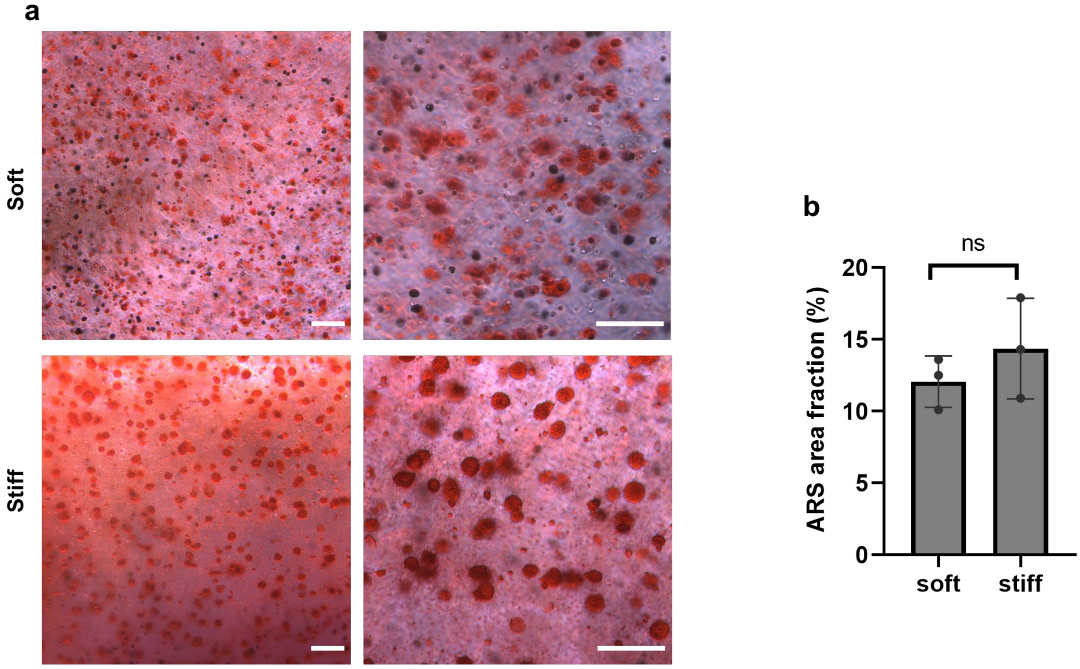


**Figure S8.** Matrix mineralization in the bioprinted constructs after osteogenic culture for 21 days. (**a**) Microscopy images of soft and stiff hydrogels stained by Alizarin Red S (ARS). Scale bars, 200 µm. (**b**) Quantification of the ARS area fraction by ImageJ. Data represented as mean ± SD. ''ns'' denotes no significant difference (n=3).

**Figure S9**

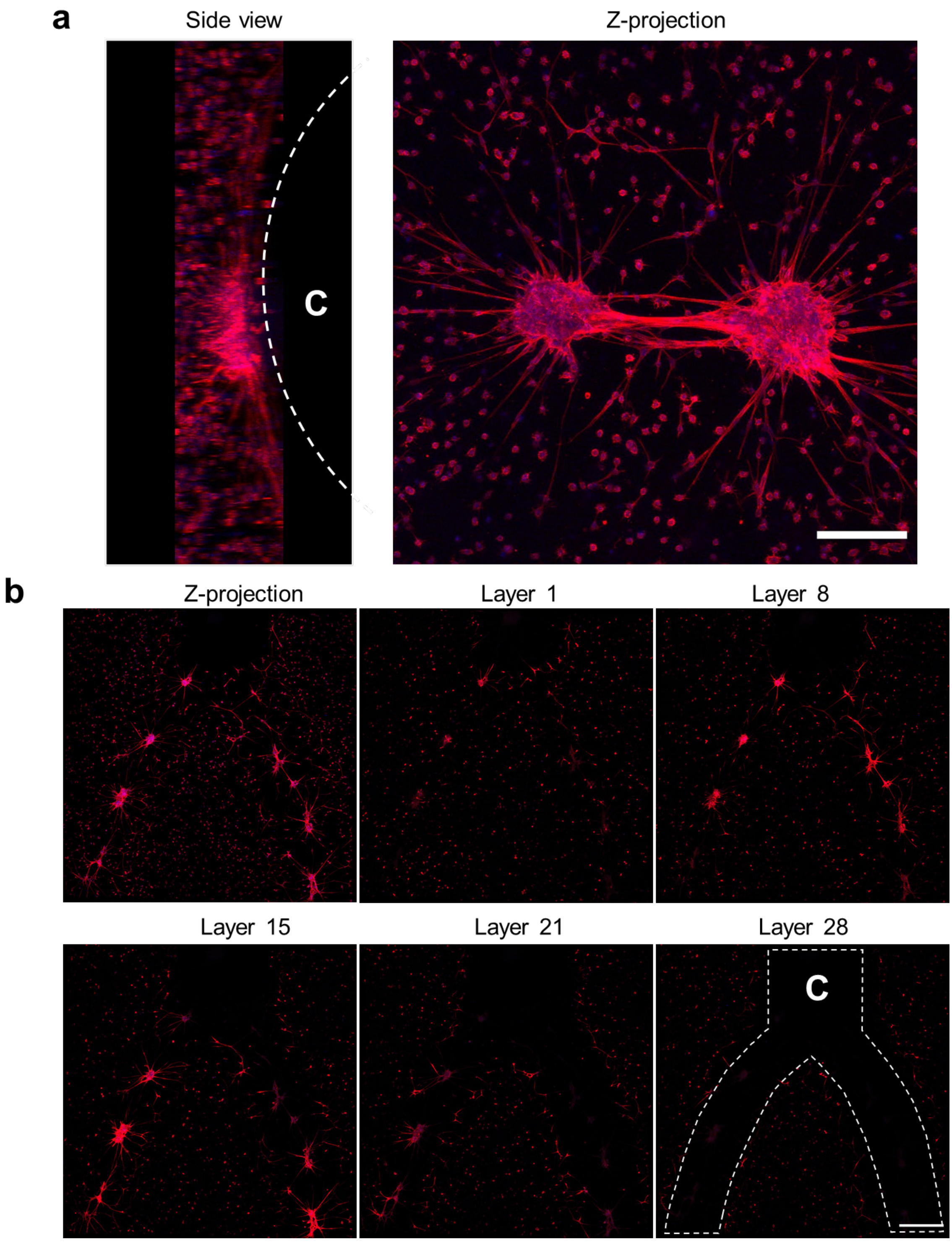


**Figure S9.** Observation of aligned cell aggregates in the channel of bioprinted constructs. (**a**) Confocal images of actin-nuclei stained hMSC cell aggregates in the big channel with side view and z-projection. (**b**) Confocal images of the cell aggregates at different depths within the printed construct. The thickness of each layer was 4.1 μm. "C" refers to the position of channels. Scale bars, 200 μm (a) and 500 μm (b).

**Figure S10**

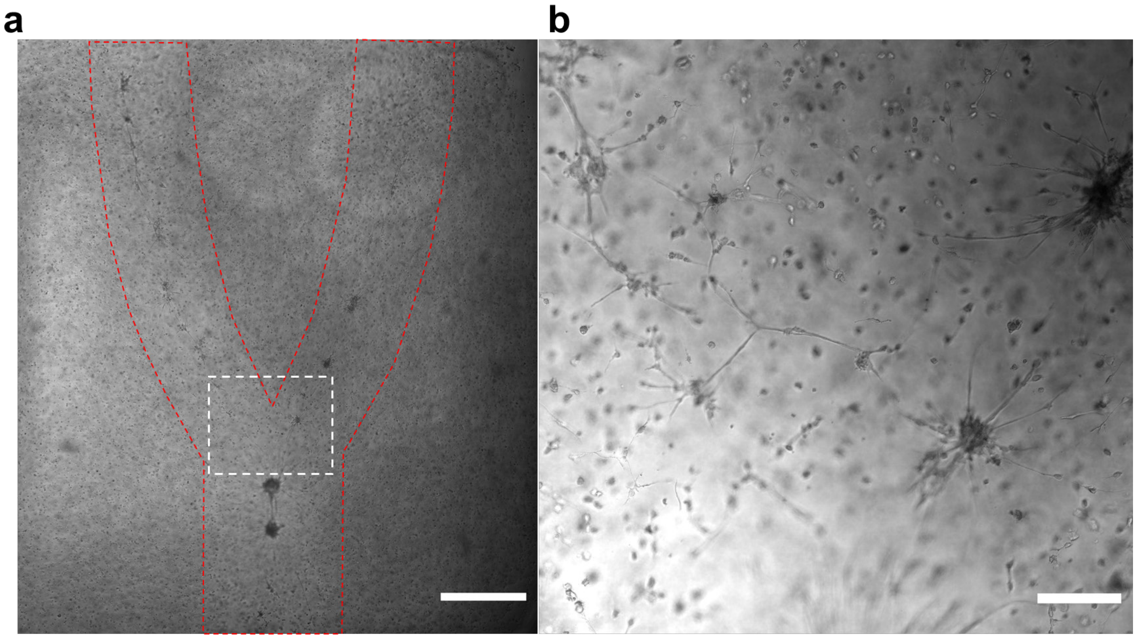


**Figure S10.** Bright-field images of the bioprinted perfusable construct (**a**) and the cell aggregates in the channel (**b**) after 7 days of osteogenic culture. Scale bars, 1 mm (a) and 200 µm (b).

**Figure S11**

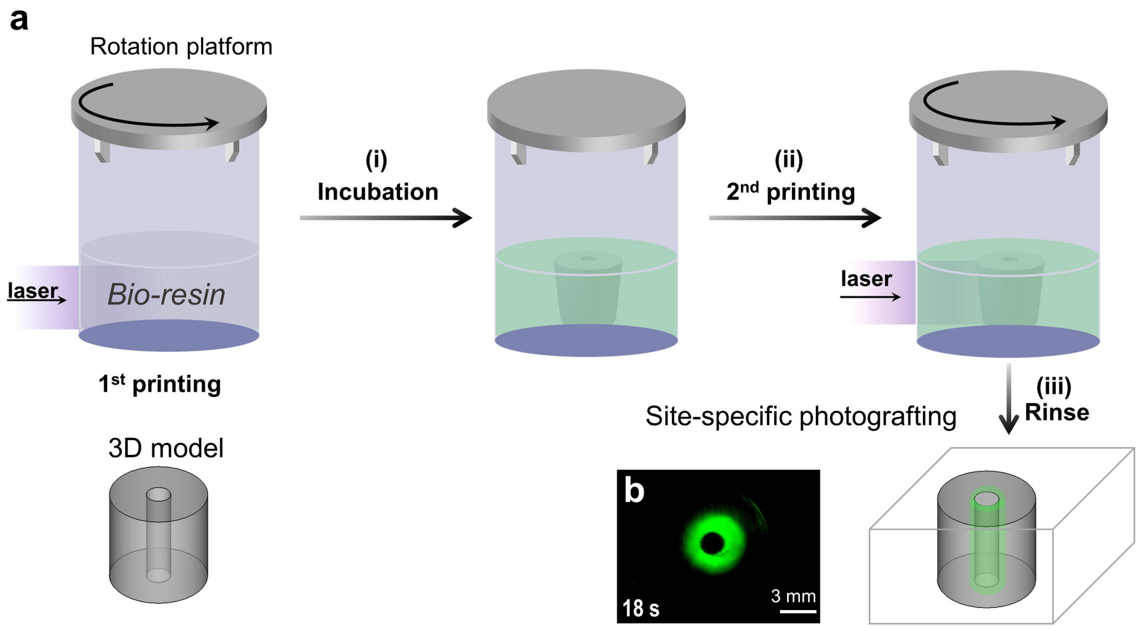


**Figure S11.** Site-specific volumetric photopatterning of chemical cues in a preformed hydrogel. (**a**) Illustration of volumetric photopatterning. (**b**) Top view of a fluorescence image of patterned channel.

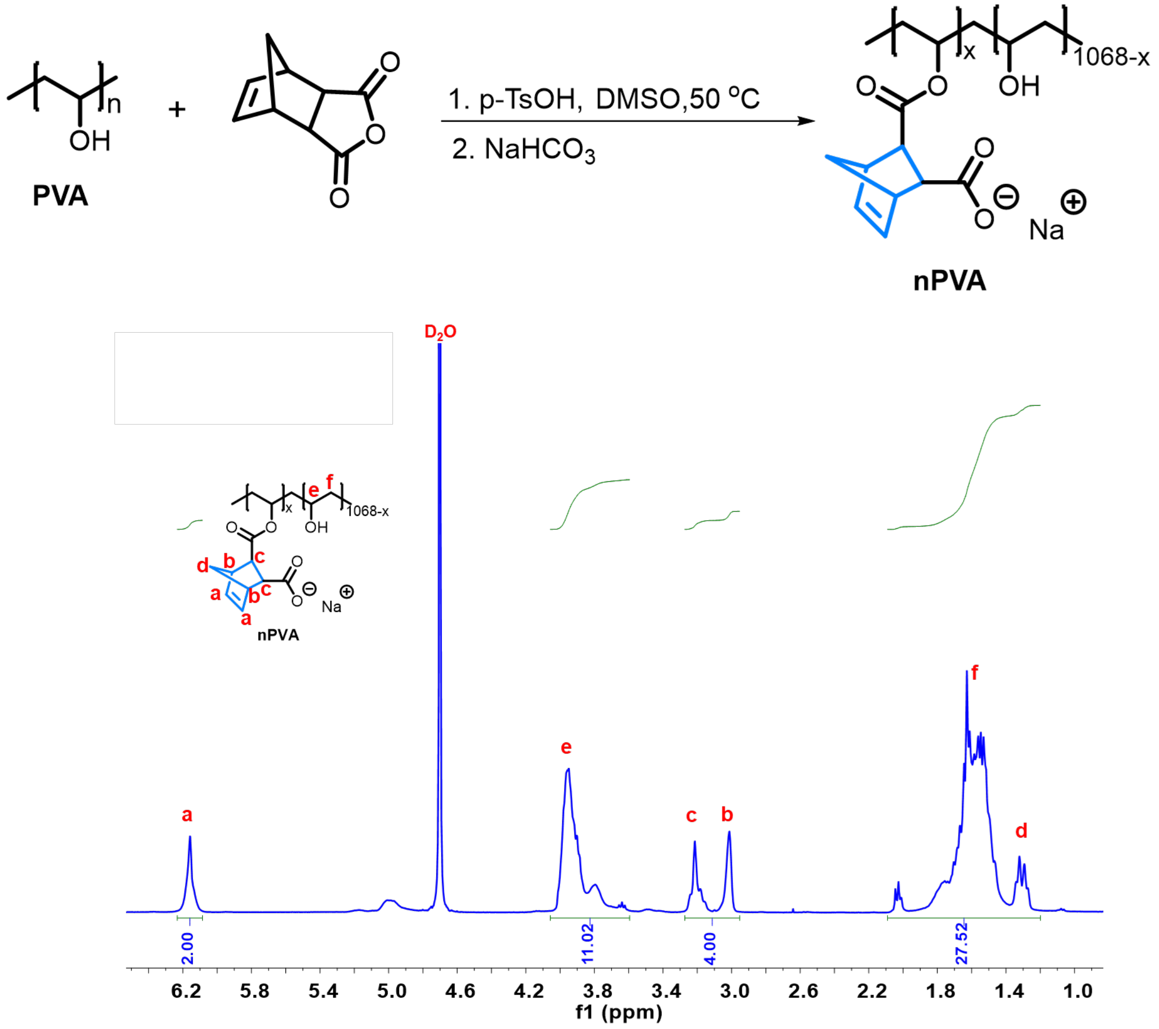


**Figure S12.** Synthetic route of nPVA and $^1$H-NMR spectrum of nPVA measured in $D_2O$. The degree of substitution (DS) was 7.8%, which was determined as described in literature.[1]

**Table S1. List of compositions of the resins in this study.**

| Entries | Resin | Polymer (% w/v) | PEG2SH (% w/v) | Gelatin (% w/v) | LAP (mM) | Pyrogallol [a)] (ppm) |
|---|---|---|---|---|---|---|
| Mix 1 | 1% nPVA | 1 | 1.1 | 5 | 2 | 50 |
| Mix 2 | 1.5% nPVA | 1.5 | 1.6 | 5 | 2 | 50 |
| Mix 3 | 2% nPVA | 2 | 2.1 | 5 | 2 | 50 |
| Mix 4 | 3% nPVA | 3 | 3.2 | 5 | 2 | 50 |
| Mix 5 | 10% GelMA | 10 | - | - | 2 | - |

[a)] Pyrogallol (50 ppm) was added to improve the stability of thiol-ene resins.

**Supplementary Videos**

**Video S1.**

**Volumetric printing procedure.** Video of the printing process of a cm-scale trabecular bone construct with a 1.5% nPVA resin. The tomographic light projection irradiated from the left, whereas the solidified component appeared at the end of the printing. Scale bar, 5 mm.

**Video S2.**

**Cell morphology in the bioprinted hydrogel constructs.** 3D illustrations of single cells in the soft and stiff hydrogel constructs after 7 days cultivation in osteogenic medium. The videos were created with IMARIS software.

**Video S3.**

**Volumetric photopatterning procedure.** Volumetric photopatterning of a helix 3D geometry within a preformed hydrogel which was incubated in a solution of 0.85 mmol $m^{-3}$ thiolated fluorescein (FITC-SH) and 1.7 mmol $m^{-3}$ LAP. Printing time, 16.3 s. Scale bar, 5 mm.
